# Oculomotor and neural correlates of semantic and morphological priming in natural sentence reading

**DOI:** 10.1101/2022.08.31.506138

**Authors:** Katarina Marjanovič, Yamil Vidal, Davide Crepaldi

## Abstract

Our current understanding of visual word identification is difficult to extend to text reading–both experiments and theories focus primarily, if not exclusively, on out–of–context individual words. Here, we try to fill this gap by studying cross–word semantic and morphological priming within sentences in natural reading, in a novel coregistration paradigm with simultaneous recording of eye movements and electroencephalography. We report results from both eye tracking measures, and, more importantly, from Fixation-Related potentials, time-locked to the fixation onset on the target word. In both, semantic facilitation clearly emerged, while we observed no effect of morphological priming. These results may indicate that morphological agreement is at least partially computed outside of the lexical-semantic system which gives rise to semantic priming. These results provide new insight into the neural correlates of semantic and morphological priming in natural reading, revealing lexical dynamics as they likely emerge in our everyday reading experience.

## 1 Introduction

Over the last decades, we have learned a lot about the mechanisms that underlie visual word identification (e.g., Adelman et al., 2010; Forster and Veres, 1998; Marelli et al., 2015; Rastle et al., 2000a; Xu and Taft, 2014). Our knowledge is so rich and articulated that it fostered the construction of fully-fledged computational models (e.g., Adelman, 2011; Coltheart et al., 2001; Davis, 2010; Perry et al., 2010). Several aspects of these models are still under debate, but our understanding has surely reached a mechanistic level, whereby theories can be entirely defined mathematically and thus produce precisely quantified predictions. This body of knowledge is very much focused on individual word identification, and unveils the fine details of the lexical dynamics that are triggered when our eyes meet letters and words.

In a somewhat parallel literature, theories of sentence reading have also been proposed that are entirely specified computationally, namely, models of eye movement during text reading (e.g., Engbert et al., 2005; Reichle et al., 2006). By design, these theories highlight the fact that the visual word identification system must engage with several words in very rapid succession (in serial models, such as E–Z Reader; e.g., Reichle et al., 2006) or even simultaneously (in parallel models, such as SWIFT; e.g., Engbert et al., 2005). This goes together with experimental work showing the large amount of words processed by the reading system when we take up connected text (from ~240 to ~300 words per minute, according to different estimates; Pelli et al., 2007; Brysbaert, 2019).

How a unique visual word identification system might deal with this issue is not obvious. Imagine a reader that is processing the sentence ‘Beth used to eat pesto every day’. When the word *eat* enters the lexical system, any model of visual word identification would predict that activation accumulates in the representation for the word eat, of course, but also in the representation of a number of similar words, orthographically (e.g., *east, feat, tea*), morphologically (e.g., *eating, ate*), and semantically (e.g., *food, dinner, pasta*)^1^. When the model gets to the following word, *pasta*, the same dynamics would get in place. If we simply generalise the workings of the word identification models, the activation triggered by the two words will sum up and get merged within the system. This is an elegantly simple solution, of course, but also one that opens up the possibility for unwarranted cross-word interactions. In the previous sentence, for example, one might mistake the word *pesto* for its higher-frequency neighbour *pasta*, which was more strongly activated by the previous word *eat*.

Of course, this risk might be compensated by other mechanisms, either within the lexicon itself or outside of it. This is the approach taken by OB1-Reader, a model of eye movement control *and* visual word identification recently proposed by Snell et al. (2018). In this model, activation does flow freely through the lexicon whatever specific word in the sentence triggered it. Excessive cross-word interference is avoided via (i) lateral inhibition between word representations and (ii) a binding system that moves any word that reaches an identification threshold from the lexicon to a spatiotopic representation of the sentence. The former mechanism is also used to ease discrimination between similar words in models of individual word identification; the latter, instead, is specific to sentence processing and OB1. This architecture is able to simulate several eye tracking effects, and does seem to be effective in limiting disruptive interference from neighbouring words in sentences; yet, it allows wide room for neighbouring words to interact within the visual word identification

This feature of OB1 licenses some experimental predictions that potentially speak to some fundamental aspects of the reading and language system. It is very well demonstrated in single word identification literature that seeing a semantically (e.g., *sell*) or morphologically (e.g., *dealer*) related word makes it easier to identify a given target (e.g., deal; Feldman, 2000; Forster et al., 1987; Grainger et al., 1991; Neely, 1977; Rastle et al.,2000a)^2^. If neighbouring words within sentences are free to interact in the lexical system, one might expect the same phenomena to occur across words during sentence reading; just as well as seeing the word *nice* in the middle of a black screen would make it quicker to identify the word *kind* in isolation, reading the former word in a sentence should make it easier to then recognize the latter word. This is the phenomenon we investigate in this paper.

Although from a rather different perspective, lexical dynamics during sentence reading were of course investigated in previous work. For example, a number of eye tracking studies have addressed semantic priming in sentences, and yielded mixed results; while some have reported facilitation (e.g., Blank and Foss, 1978; Van Petten et al., 1997), others have found that lexical priming is easily overridden by sentence-level factors, such as discourse context and predictability (e.g., Duffy et al., 1989; Morris, 1994; Morris and Folk, 1998; Traxler et al., 2000). For example, Morris (1994) reported savings in the identification of a target word (e.g., *moustache*) from related word primes (*barber* and *trimmed*) in sentences like ‘the gardener talked as the barber trimmed the moustache’. However, this effect dissappered in sentences, in which the second noun (i.e., *gardner*, in the above example) become the agent of the priming verb (e.g., ‘the gardener talked to the barber and trimmed the moustache’) and distorted the semantic connection at the sentence level (gardeners do not trim moustache, typically), suggesting that the facilitation on the target word was not primarily lexical, but more dependent on the sentence-level context.

Morphological effects were also heavily studied in sentence reading (e.g., Barber and Carreiras, 2005; Kos et al., 2010; Kutas and Hillyard, 1980, 1984a; Weber and Lavric, 2008). Most of this work, however, is rather difficult to interpret in terms of cross–word priming, given that it is based on morpho– syntactic violations. For example,Bates et al. (1996) report quicker repetition in noun phrases with a correct agreement for gender (e.g., *‘brutta casa’, ugly_F_ house_F_*) vs. gender-mismatch controls (*\*‘brutto casa’, ugly_M_ house_F_*). These effects are typically interpreted in terms of costs related to the morphological violation. One could surely take the opposite perspective and imagine that a morphological match yields time savings, but a link to classic priming data would remain problematic as the ‘control’ condition is typically neutral in standard priming, while it entails a frank morpho-syntactic violation here. The very concept of morphological (and semantic, for what matters) violation is ruled out in individual word paradigms. Also, many of these experiments involved rather minimalistic environments such as word pairs (e.g., Goodman et al., 1981; Samar and Berent, 1986; West and Stanovich, 1988); it is not obvious, then, whether these data would generalise to more naturalistic stimuli, like sentences or entire passages of text.

In the present study, we make use of a paradigm that overcomes the limitations illustrated above. We simply asked participants to read sentences for comprehension, and checked for cross-word priming as shorter fixations on words that were preceded by semantically or morphologically congruent words. This approach involves more naturalistic stimuli, avoids any morphological or semantic violation, and allows us to use the material that is typically adopted in the single word literature; content words that are semantically and/or morphologically congruent are located close to each other in a sentence (e.g., ‘… forks and spoons…’), so that if the relevant information persists in the lexical system, we should observe savings in the identification time (that is, fixation duration) of the latter word. Quite conveniently, eye tracking would also allow us to estimate the time course of the eventual effects, through a comparison between earlier fixation measures (e.g., first–of–many fixation duration) and later eye movement metrics (e.g., gaze duration).

As far as semantics goes, we are aware of only two eye-tracking studies that adopted a similar paradigm (in addition to Morris (1994), which we described above).Carroll and Slowiaczek (1986) focused on how the structure of the sentence would affect lexical processing. They found that priming is influenced by the syntactic structure of the sentence, so that it is observed only when the semantically related prime and target appear in the same clause (e.g., ‘The guard saluted *the king* and *the queen* in the carriage, but they didn’t notice’). One issue with this experiment, however, is the lack of control over target word predictability. As observed in, e.g., Otten and Van Berkum (2008), there is widespread prediction of the upcoming word during sentence reading, which makes it difficult to unambiguously attribute Carroll’s and Slowiaczek’s results to cross–word priming. Facilitation may actually come from the on-line prediction of the target, to which the prime surely contributes as part of the sentence, but which it doesn’t determine per se. The interaction between lexical representations in the mental lexicon may not be the driving force behind these results.

A similar issue affects the study of Camblin et al. (2007). These authors did gather data on target word predictability, but mean cloze probability in their semantically congruent condition was very high (averaging .36). Thus, on–line prediction based on sentence context may have played a major role in this experiment too, making unclear the contribution of lexical dynamics.

Things are not entirely clear on the morphological side either. For example, Paterson et al. (2011) investigated how the prior exposure to morphologically related words may influence target word processing. Their prime–target pairs were either semantically transparent (e.g., *marshy-marsh*), had only an apparent morphological relationship (e.g., *secretary-secret*), or were morphologically unrelated but as orthographically similar as in the previous conditions (e.g., *extract-extra*). Priming effects were observed in the semantically transparent pairs, but were absent in the remaining two conditions. This study clearly shows that the observed morphological priming effect is not driven by the prime–target (morpho-)orthographic relationship (e.g., Rastle et al., 2004). However, as the authors themselves acknowledge, it is impossible to establish whether the observed effect is morphological in nature or is rather a more general semantic effect, of the sort we would find with words like *cat* and *dog*— without such a semantic control condition, this question cannot be addressed.

Semantic and morphological processing during sentence reading has been widely explored also through scalp electrophysiology (EEG), particularly through Event-Related Potentials (ERP) where sentences were presented one word at a time. In this literature, semantic processing is typically linked to the N400 ERP component, which denotes a negative-going deflection starting around 250ms and peaking around 400ms after stimulus onset, with a centro-posterior distribution (e.g., Kutas and Federmeier, 2000, 2011; Federmeier, 2007; Hagoort, 2003; Traxler and Gernsbacher, 2011). In semantic priming, N400 is larger (i.e., more negative-going), when the target word is preceded by semantically unrelated word, and smaller (i.e., more positive-going) when the target word is anticipated by a semantically related word (e.g., Kutas and Hillyard, 1984b; Rugg, 1985).

The N400 literature is enormous (see, e.g., Kutas and Federmeier, 2011, for an overview), but its interpretation remains somewhat controversial. Traditionally, it has been interpreted either as an index of facilitated lexical access (e.g., Lau et al., 2009; Rugg, 1990) or as an index of access to the conceptual knowledge of a word (e.g., Federmeier, 2007; Kutas and Federmeier, 2000); these interpretations are quite close to the classic accounts of lexical and semantic priming in the identification of isolated words. However, the N400 has been also associated with postlexical processes, such as semantic context integration (e.g., Brown and Hagoort, 1993; Holcomb, 1993). It remains indisputable, however, that the N400 provides information about the time course of the semantic processing; in this context, its onset can be interpreted as an upper time limit of the initial access to the word meaning (Dimigen et al., 2011).

The N400 has also been linked to morphological processing. In priming paradigms, the N400 is generally larger when the target word is preceded by a morphologically unrelated prime, and smaller when it is preceded by a morphologically related word (e.g., Holcomb and Grainger, 2006). This modulation has been consistently reported with stem (e.g., *hunter-HUNT*) and repetition priming (e.g., *hunt-HUNT*), both in masked (e.g., Holcomb and Grainger, 2006; Morris et al., 2008; Lavric et al., 2007) and overt priming (e.g., Lavric et al., 2010; Smolka et al., 2015). In some studies, morphological priming was also reported as related to an earlier ERP component, sometimes dubbed N250, which some scholars have interpreted as a pre-lexical stage of morphological analysis, one that is strongly informed by orthography (e.g., acting also on pseudo-derived words like *corner* or *irony*; Morris et al., 2008; Morris and Stockall, 2012), as opposed to a more semantically-informed, post-lexical morphological level (see, e.g., Crepaldi et al., 2010; Taft, 2004, for models implementing this distinction).

The evidence is even less consistent when it comes to inflectional morphological priming, which is most commonly addressed in the comparison between regular and irregular English verbs. Some studies observed an N400 modulation with regular, but not irregular verbs (e.g., Rodriguez-Fornells et al., 2002), while others did not find this regularity effect (Justus et al., 2011, in an auditory priming paradigm). Other studies observed the biphasic N250/N400 modulation using masked priming, but only with regular verbs (e.g., Rastle et al., 2015), while others, again, observed the same pattern of results with both regular and irregular verbs (Morris and Stockall, 2012).

Somewhat similarly to the eye tracking literature illustrated above, most of these studies obtained their results with either words in isolation or sentence-based paradigms that differ from the natural reading experience in a number of ways (e.g., Degno et al., 2018; Dimigen et al., 2011; Metzner et al., 2017). In Rapid Serial Visual Presentation (RSVP) paradigms, the words making up a sentence are presented in isolation in rapid succession in the middle of a screen, while participants are instructed to maintain their gaze still and avoid blinking; this serves the purpose of having a clear time onset for the target word, from which ERPs are measured. However, this setup is quite unnatural, and is likely to bring additional cognitive load, as the participants try to follow the given instruction and become aware of their otherwise unconscious eye movements. This could potentially have uncontrolled effects on the observed neural signatures. Further, each word is presented on a screen for a predetermined amount of time (typically around 400ms), which is significantly longer than the average time readers spend on a fixated word (200-250ms; e.g., Sereno and Rayner, 2003). This slows down the reading speed, which could in turn have unwanted effects on the processing speed (Dimigen et al., 2011). Another important difference lies in how each individual word is visually processed to enter the reading system. In RSVP paradigms, the reader is forced to read each word, and in a strictly serial order. This contrasts with normal reading, during which readers freely determine not only how long each word will be fixated, but also which word will be fixated next. We do not necessarily read in a strictly serial manner; some words are fixated more than once, while others are skipped, and regressive saccades towards previously fixated words are quite frequent (e.g., Dimigen et al., 2011). Finally, RSVP paradigms do not allow for the preprocessing of the upcoming words in parafoveal vision, which is of course widespread in normal reading (e.g., Degno et al., 2018; Dimigen et al., 2011).

To heal these limitations, a novel paradigm was introduced recently, which is based on the simultaneous recording of EEG and eye movements (e.g., Dimigen et al., 2011, 2012; Degno et al., 2018). Co-registration allows to time-lock the EEG signal to specific eye movements, such as a fixations on the target word; therefore, these event-related potentials are referred to as Fixation-Related Potentials (FRPs). In this paradigm, participants are presented with whole sentences (or passages of text, in principle) and are allowed to freely move their eyes as they process them (Degno et al., 2018).

We are aware of no co-registration study that has addressed morphological priming in natural sentence reading. However, some evidence is available on semantic processing/priming. In an experiment where participants were asked to read lists of words (e.g., *kitchen forest tree blur drive*), Dimigen et al. (2012) found an effect in the FRPs that might be described as semantic priming. Indeed, the data showed that unrelated, related and identical words at position n (which can be interpreted as primes) elicit large, intermediate, and small N400 amplitudes, respectively, on words at position n+1 (which can be interpreted as targets; Dimigen et al., 2012). This FRP N400 component had a similar spatial distribution as the corresponding ERP, but emerged substantially earlier (as early as 80ms after the onset of the fixation on word n+1). In another study that used actual sentence reading, Kretzschmar et al. (2009) found somewhat different results. Their design focused on disentangling predictability from semantic relatedness *per se*. In the critical comparison between related and unrelated words, both unpredictable in the sentence, they found no significant difference in the N400 time window. Note, however, that – contrary to Dimigen et al. (2012), the single word priming literature more generally, and the present experiment – the manipulation here involved the word n+1 (the target), while the word n (the prime) remained fixed.

To sum up, building on previous previous work, we devised a paradigm that unambiguously assesses semantic and morphological cross–word priming during sentence reading. Primes and targets are embedded into sentences and put close together in a coordinating phrase (e.g., ‘Paul entered a room with *a table* and *a chair*, which didn’t really look like a kitchen’). Their semantic (S) and morphological (M) relationship is then independently manipulated, e.g., ‘a table and a chair’ (S+M+) vs. ‘a dog and a chair’ (S-M+) vs. ‘some tables and a chair’ (S+M−) vs. ‘some dogs and a chair’ (S-M-). In the eye tracking data, priming is taken to occur if fixations on the target word (*chair*) are shorter after semantically and/or morphologically congruent primes. In the EEG data, priming is taken to occur if FRPs locked to the target words elicit a more positive- or negative-going deflection after semantically and/or morphologically congruent primes.

This paradigm, as illustrated above based on English, has two main problems though. First, as already pointed out by Paterson et al. (2011), morphological relationship brings about orthographic relationship (plural words share a final –s), which makes it difficult to disentangle these two effects, in particular for shorter words. Second, the presence of determiners in the critical bit of the sentence is not ideal, for at least two reasons. Although articles are typically skipped during reading (e.g., Angele and Rayner, 2013), they could potentially attract at least some fixations, which would be difficult to handle—should they count as fixations on the target word? Or perhaps they would determine quite some more skipping of the target word itself? Also, and probably more relevant, articles contain morphological information, which would be extracted by the readers (foveally or parafoveally), thus blurring the whole picture—would morphological priming come from the content words or the determiners, or some cross–talk between the two? How does this affect the pattern of results, if it does at all? How would then results be comparable to those emerging from the individual word literature?

Here is where Slovenian, the language that we used for this experiment, turns out to be handy. Slovenian does not use determiners, so that primes and targets would sit alone in the critical coordinating phrase (e.g., *‘miza in stol’*, ‘(a) table and (a) chair’). Also, Slovenian is inflectionally very rich—it has 6 different cases, 3 different genders, and 3 different grammatical numbers, with noun declension introducing distinct suffixes for different combinations of the three. Nouns can thus be inflected in the same way (i.e., in number and case), but still have orthographically different suffixes (e.g,. *‘avtomobil-i’*, *cars*, plural, nominative; and *‘učiteljic-e’*, *teachers*, plural, nominative). This peculiarity of the Slovenian language thus enables us to rule out any orthographic contribution to morphological priming.

## 2 Methods

### 2.1 Participants

41 right handed, native Slovenian speakers (F=24) took part in the study. Their mean age and education was 26.29 (range=19–46) and 15.66 years (range=12–20), respectively. They all grew up in a monolingual environment and had normal or corrected-to-normal vision. They all provided their informed consent to take part in the study before the beginning of the experiment, and received a 25 Euro compensation. Due to unsuccessful calibration of the eye tracking system, 2 participants were then excluded from the analysis.

### 2.2 Materials

The stimuli set comprised 40 sentences, in which two nouns appeared one after the other, separated by the conjunction in, and (e.g., *‘kolesar ni bil pozoren na **avto** in **tovornjak** in je zato povzročil nesrečo’*; ‘the cyclist was not paying attention to **a car** and **a truck** and therefore caused an accident’). In our design, the first noun (‘avto’, car, in the example) is the prime word, while the second noun (*‘tovornjak’*, *truck*, in the example) is the target word.

These sentences appeared in 4 different conditions, where primes and targets were (i) related in meaning, and inflected in the same grammatical number (*‘avto–tovornjak’, (a) car–(a) truck*); (ii) related in meaning, but not inflected in the same grammatical number (*‘avte–tovornjak’, (some) cars–(a) truck);* (iii) unrelated in meaning, and inflected in the same grammatical number (‘*lužo–tovornjak’*, (*a) puddle–(a) truck);* (iv) unrelated in meaning, and not inflected in the same grammatical number (*‘luže–tovornjak’*, (*some) puddles–(a) truck*). Both the carrier sentences and the target words were kept identical across conditions; only the prime varied, to determine morphological and/or semantic relatedness in a crossed, 2-by–2 design.

While constructing the sentences, we took a series of measures to guarantee a fair assessment of cross–word priming. First, the prime and the target word always appeared within the same syntactic clause (Carroll and Slowiaczek, 1986; Morris and Folk, 1998). Second, as anticipated above, primes and targets never shared the same orthographic suffix, and only shared the same final letter in 9 cases (5.6% of the stimuli set), so as to rule out any substantial contribution from form priming. As mentioned above, this is easily obtained in Slovenian through the use of prime and target nouns with different gender (e.g., *‘brisač-i’, towels*, and *‘ležalnik-a’*, *deckchairs*, are both dual nouns in the nominative case). Furthermore, target words never appeared in either clause or sentence final position, and were never followed by a comma—these conditions may in fact elicit wrap-up effects, with longer fixations depending on syntactic and/or semantic integration (Warren et al., 2009). Finally, we also controlled the position of the prime and the target word on the screen—they never appeared as the first or the last words in a line.

Primes and target features are illustrated in Table 1. They were matched as closely as possible for length and frequency. This latter was taken from the Slovenian corpus [Gigafida](https://viri.cjvt.si/gigafida/). Sentences were 12 to 20 words long (mean=15.6), and included 63 to 138 characters overall (mean=92.3). The prime words came 19 to 62 characters into the sentence (mean=36.7). Twenty-five sentences were displayed in two lines of text, whereas 15 occupied three lines of text on the screen.

**Table 1:** Frequency and length of our stimuli across conditions. We report means and SDs. Frequency is taken from the Slovenian corpus *Gigafida*, log transformed, and based on word form.

|  | **S+M+**<br>**prime** | **S+M-**<br>**prime** | **S-M+**<br>**prime** | **S-M-**<br>**prime** | **Target** |
| --- | --- | --- | --- | --- | --- |
| Frequency | 1.46 (0.54) | 1.35 (0.53) | 1.34 (0.58) | 1.36 (0.52) | 1.32 (0.50) |
| Length | 6.60 (2.10) | 6.57 (2.09) | 6.97 (2.11) | 7.25 (2.32) | 6.57 (2.04) |

S+/-: semantically congruent/incongruent; M+/-: morphologically congruent/incongruent.

#### Predictability

In order to unambiguously assess the cross-word priming effects and also to prevent an excessive skipping of the target words, we made sure that these latter were not too predictable. A cloze probability task was set up (Kutas and Hillyard, 1984b) with all the sentences that were then used in the experiment proper. A separate sample of 80 participants (F=54; mean age=32.38; age range=20–65), none of whom took part in the main experiment, were presented with the experimental sentences up to the pre–target word and were asked to complete them with the first word that came to mind. Because of our design, each target was anticipated by four different primes, in four otherwise identical sentences (e.g., *‘Kolesar ni bil pozoren na **avto/avte/lužo/luže** in **tovornjak***…’). To make sure that target predictability was similar (and low) across conditions, all of these four sentences were tested in the Cloze probability task, using a Latin Square design (each participant was presented with only one item in each sentence quadruplet, rotated across conditions).

In the final set of stimuli, no target word had a cloze probability higher than .2. Means and SDs across conditions are illustrated in Table 2. Some form of semantic (but not morphological) priming emerges here as a slightly higher CP in the semantically congruent (S+) conditions—participants tended to guess the word that would be used in the stimulus sentence more often after the semantically related primes. This effect is likely unavoidable, but, critically, very small in our set of items (see Table 2). However, to be entirely sure that this slight difference in CP was not a critical player in the experiment, we reran any eye tracking analysis that would reveal semantic priming after excluding sentences with *CP* > .10.

**Table 2:** Mean (SD) cloze probability across conditions.

|  | S+M+ | S+M- | S-M+ | S-M- |
| --- | --- | --- | --- | --- |
| Cloze probability | .05 (.05) | .06 (.07) | .01 (.03) | .01 (.03) |

S+/-: semantically congruent/incongruent; M+/-: morphologically congruent/incongruent.

#### Semantic relatedness

We also tested the strength of the semantic relatedness between the target words and their primes. A further separate sample of 21 participants (F=12; mean age=38; age range=25–58), none of whom took part in the eye tracking experiment or the cloze probability task, was asked to rate each prime–target pair for similarity in meaning on a 1-to-5 scale (1, not similar at all; 5, very similar). Because, again, each target was associated with two different primes (primes were tested only in one morphological form here), participants were rotated over conditions (related vs. unrelated) in a Latin Square design. The results of this pre–test are illustrated in Figure 1, and show that semantically congruent primes were rated as substantially more related to their targets than semantically incongruent ones, consistently across targets (mean and SD are 3.38 and 0.54, respectively, for the congruent condition; and 1.74 and 0.42 for the incongruent condition).

**Figure 1:**
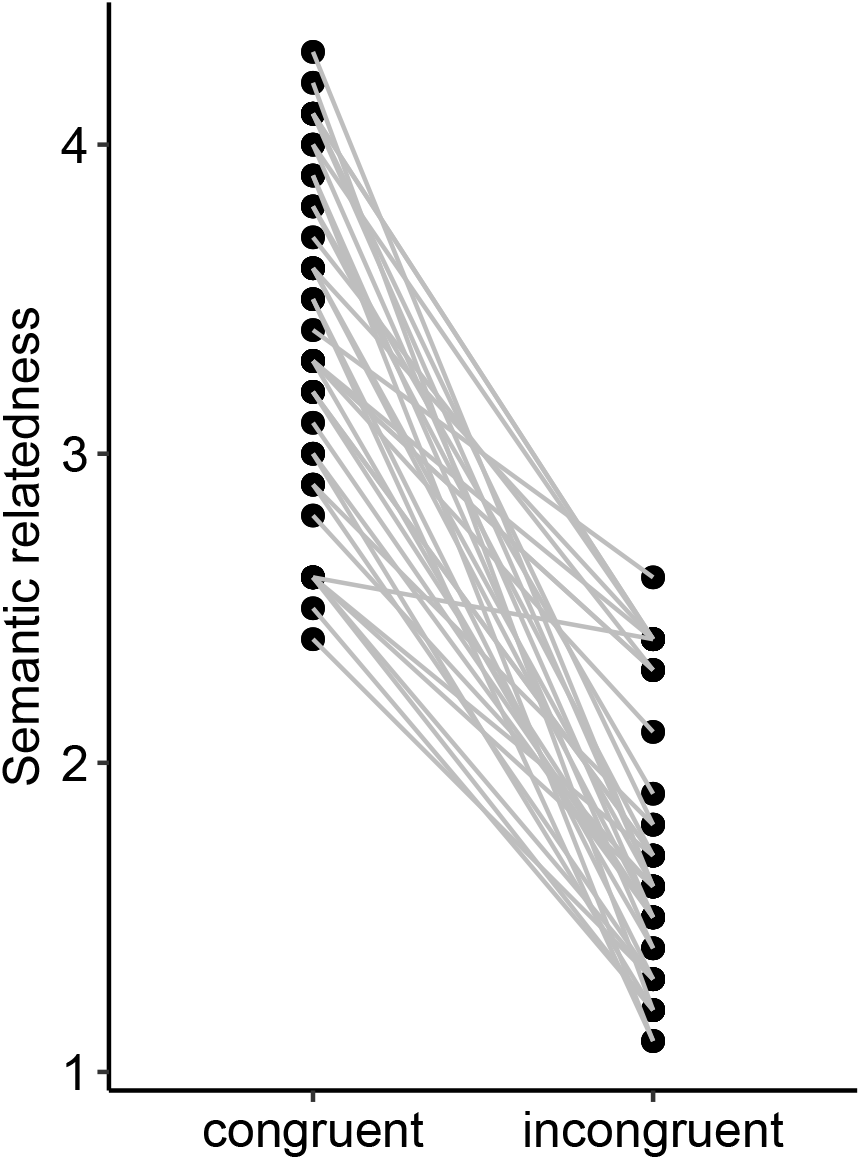
Semantic relatedness in our Congruent (left) and Incongruent (right) pairs. Grey lines connect corresponding primes, those that were used with the same target.

### 2.3 Procedure

The initial calibration of the eye-tracking system consisted of a standard 9-point grid. During the experiment, the calibration was automatically checked with a fixation point before the beginning of each trial, and was repeated when necessary.

Participants were instructed to silently read the sentences at their own pace. On roughly 30% of the trials, sentence reading was followed by a 2AFC comprehension question, to ensure that participants were actually reading and understanding the sentences.

The trial timeline was as follows. A fixation cross appeared on the screen, exactly at the location where the first letter of the sentence later appeared. Sentence presentation was triggered by participants’ fixation of the cross. The sentence was then displayed, and remained visible until the participants fixated another cross, located at the bottom of the screen. This led to the presentation of either the drift correction fixation point of the following trial, or the comprehension question.

In order to reduce eventual repetition effects, each participant was presented with each sentence in only two out of the four experimental conditions, in a standard a Latin Square design. Each session was divided in two blocks, and two similar sentences never appeared in the same block. To further minimize repetition effects, and also to reduce participants’ awareness of the goal and the structure of the experiment, each block also included 60 filler sentences, which had a different structure and were not analyzed (e.g., *‘Nina je v smetnjaku našla mlado mucko in se odločila, da jo posvoji.’*; ‘Nina found a kitten in a trash can and decided to adopt it.’). The critical sentences were arranged across blocks in such a way that each block included 10 sentences per condition. Overall, each participant read a total of 200 sentences, in two separate blocks of 100 each (i.e., 40 relevant and 60 filler sentences). The experiment lasted about 50 minutes.

### 2.4 Apparatus

The experiment was designed and ran in `Experiment builder` (SR Research Ltd., Kanata, ON, Canada). Participants sat 56 cm from the computer screen where the stimuli were displayed. Their head was stabilized through a chin rests. Eye movements were recorded with an `SR Eyelink 1000+`, at a sampling rate of 1000 Hz, from the dominant eye only. The EEG signal was recorded with a standard 64-channel, 10-20 montage (`BioSemi ActiveTwo`) system, at a sampling rate of 512 Hz. Additionally, 4 EOG channels were used to record the EEG signal related to eye movements.

### 2.5 Coregistration of eye movements and EEG signal

Following the procedure described in (Degno et al., 2018), the stimulus display computer, through the parallel port, sent a message to the computer recording the eye movements, and a trigger to the computer recording the EEG signal, to mark the beginning and the end of the experiment and of each trial. These triggers were used to establish the synchronization of the two recordings offline, which was done in `MATLAB`, with the `EYE-EEG` extension (Dimigen et al.,2011) of the `EEGLAB` toolbox (Delorme and Makeig, 2004).

### 2.6 Eye movement data preprocessing and analysis

Two interest areas were created for each sentence around the prime and the target words, using `Data Viewer` (SR Research Ltd., Kanata, ON, Canada). Trials with target word skips (3.76%) were discarded. Further, only fixations that occurred during first-pass reading were included in the analysis.

Because we were interested in exploring the time course of the eventual cross-word priming, we analyzed a number of eye tracking metrics, namely, first-of-many fixation duration^3^, gaze duration and total looking time. Moreover, in order to explore the possibility of parafoveal-on-foveal priming, we also analyzed target skipping and gaze duration on the primes.

Data were analyzed with `R` (R Development Core Team, 2008), using `Rstudio` (RStudioTeam, 2016) and the `lme4` package (Bates et al., 2015) for fitting (generalised) linear mixed models. Model estimates and effect sizes were obtained using the package `Effects` (Fox and Hong, 2009). Continuous dependent variables were modelled as a function of semantic and syntactic congruency, with subjects and target words as random intercepts. Statistical significance was checked both for model parameters and for predictors overall. Effects were checked for their dependence on outliers following Baayen (2008)—models were re–run after excluding data points whose standardised residuals were larger than 2.5 in absolute value. There were no effects that would be significant only with (or without) outliers; the data reported here refer to the results of the models including all data points.

### 2.7 EEG data preprocessing

The EEG data preprocessing was performed in `MATLAB` (Version: 9.3.0.713579 (R2017b); MathWorks, Inc., Natick, MA, USA), mainly through the `EEGLAB` (Version 14.1.1) (Delorme and Makeig, 2004) toolbox and custom code. Initially, the data were band-pass filtered with a high-pass filter of .1 Hz and a low-pass filter of 30 Hz (Hamming windowed sinc FIR filter). Eye movement data were imported and synchronized using the `EEGLAB` extension `EYE-EEG` (Dimigen et al., 2011). Continuous data was then segmented. The segments were time-locked to fixation onsets on the target words and included 100ms before and 500ms after fixation onset. Noisy channels were rejected, using the `EEGLAB`’s `pop rejchan` function (Delorme and Makeig, 2004) using three different methods: (1) Kurtosis treshold (set to 4σ); (2) joint probability treshold (set to 4σ); and (3) abnormal spectra (checked between 1 and 30 Hz, with a threshold of 3σ). Trials with extreme values (±300*μV*) were rejected. Next, we applied Principal Component Analysis (PCA) to reduce the dimensionality of the data by retaining the first 24 principal components, and decompose the data using Independent Component Analysis (Extended Infomax). Independent components associated with ocular artifacts were identified and rejected using the `EYE-EEG` extension (Dimigen et al., 2011). `EYE-EEG` picked independent components that shared temporal covariance higher than .8 with eye movements, and marked them as oculomotor artifacts (Plöchl et al., 2012). Further, trials were baseline-corrected to 200ms before the fixation onset, and trials containing extreme values (±200 *μV*) and improbable trials (`EEGLAB pop_jointprob` 4σ for both single channel and all channels) were rejected^4^. Data were then re-referenced to the average of all the scalp electrodes. Missing channels were interpolated (EEGLAB `pop_interp`, ‘spherical’). Finally, trials were divided into conditions and averaged within participant.

### 2.8 EEG data analysis

EEG data statistical testing was performed through a nonparametric clustering method (Bullmore et al., 1999), as implemented in the `Fieldtrip lite` (Oostenveld et al., 2011) plug-in for `EEGLAB` (Delorme and Makeig, 2004). Included were all scalp electrodes and all time points for a time window of 500ms, starting at the fixation onset. The clustering method was applied as follows. First, every time point of every channel of the conditions to contrast were statistically compared (we used a nonparametric permutation t test with 5000 permutations). Next, the temporally and spatially adjacent points with p values < .05 were used to form candidate clusters. For each of this candidates, a cluster-level statistic was calculated by summing the t values within each cluster. The significance of the candidate clusters was assessed via nonparametric permutation test, in which the conditions were randomly shuffled and cluster-level t values were calculated in the same manner as before. This step was repeated 5000 times, and on each iteration, the most extreme cluster-level t value was used to create a null distribution. The significance of the observed candidate clusters was then calculated as the proportion of expected t values under the null hypothesis that were more extreme than the observed ones.

## 3 Results

### 3.1 Behavioral results

All participants responded correctly to at least 89% of the comprehension questions (overall mean=95%, SD=2.15%), which suggests that they understood the sentences well and performed the task appropriately.

### 3.2 Eye movement results

The overall descriptive statistics for the variables that we considered in the analyses are reported in Table 3. Spearman’s rho correlation coefficient was used to assess the relationship between gaze duration (GD) and total looking time (TLT). There was a fairly strong correlation between the two (*r_s_* = .577), which reflects the proportion of refixation.

**Table 3:** Means (and standard deviations) across conditions for the eye–tracking metrics that we considered in this paper. Statistics are reported in ms, and are based on unaggregated data. Note: Prime, prime fixation duration (which tracks target–on–prime effects); FoM, first–of–many fixation duration; GD, gaze duration; TLT, total looking time; S+/−, semantically congruent/incongruent; M+/-, morphologically congruent/incongruent.

|  | S+M+ | S+M- | S-M+ | S-M- |
| --- | --- | --- | --- | --- |
| Prime | 231 (146) | 230 (146) | 241 (154) | 251 (153) |
| FoM | 160 (97) | 155 (86) | 168 (101) | 169 (107) |
| GD | 248 (136) | 255 (149) | 273 (159) | 275 (168) |
| TLT | 395 (233) | 421 (249) | 503 (344) | 516 (371) |

We firstly assessed priming based on parafoveal information in the form of a target-on-prime effect—essentially, we checked whether the overall duration of all first-pass fixations on primes were shorter when the following target words were semantically and/or morphologically related^5^. This phenomenon would belong to the class of the hotly debated Parafoveal-on-Foveal (PoF) effects (e.g., Just and Carpenter, 1983;Henderson and Ferreira, 1993;Inhoff et al., 2000). Our data suggests a significant semantic priming effect, *F*(1,2968) = 4.151, *p* = .043 (corresponding model parameter, *t*(2968) = 2.038), but no morphological priming, *F*(1,2968) = .333, *p* = .565, nor interaction between semantic and morphological congruity, *F*(1,2968) = .531, *p* = .467. The estimated effect size was 16.1 ms for semantic priming and 4.6 ms for morphological priming.

First-of-many (FoM) fixation durations also revealed a significant effect of semantic congruity, *F*(1,3000) = 8.843, *p* = .003 (corresponding model parameter, *t*(3000) = 2.974), but no morphological priming, *F*(1, 3000) = .136, *p* = .712, nor interaction between semantic and morphological congruity, *F*(1, 3000) = .636, *p* = .426. The estimated effect size was 11.09 ms for semantic priming and 1.36 ms for morphological priming. Figure 2a presents the model-based estimates for FoM. This pattern did not change when we considered only those trials in which the primes were fixated (semantic effect: *F*(1,2861) = 4.77, *p* = .029).

**Figure 2:**
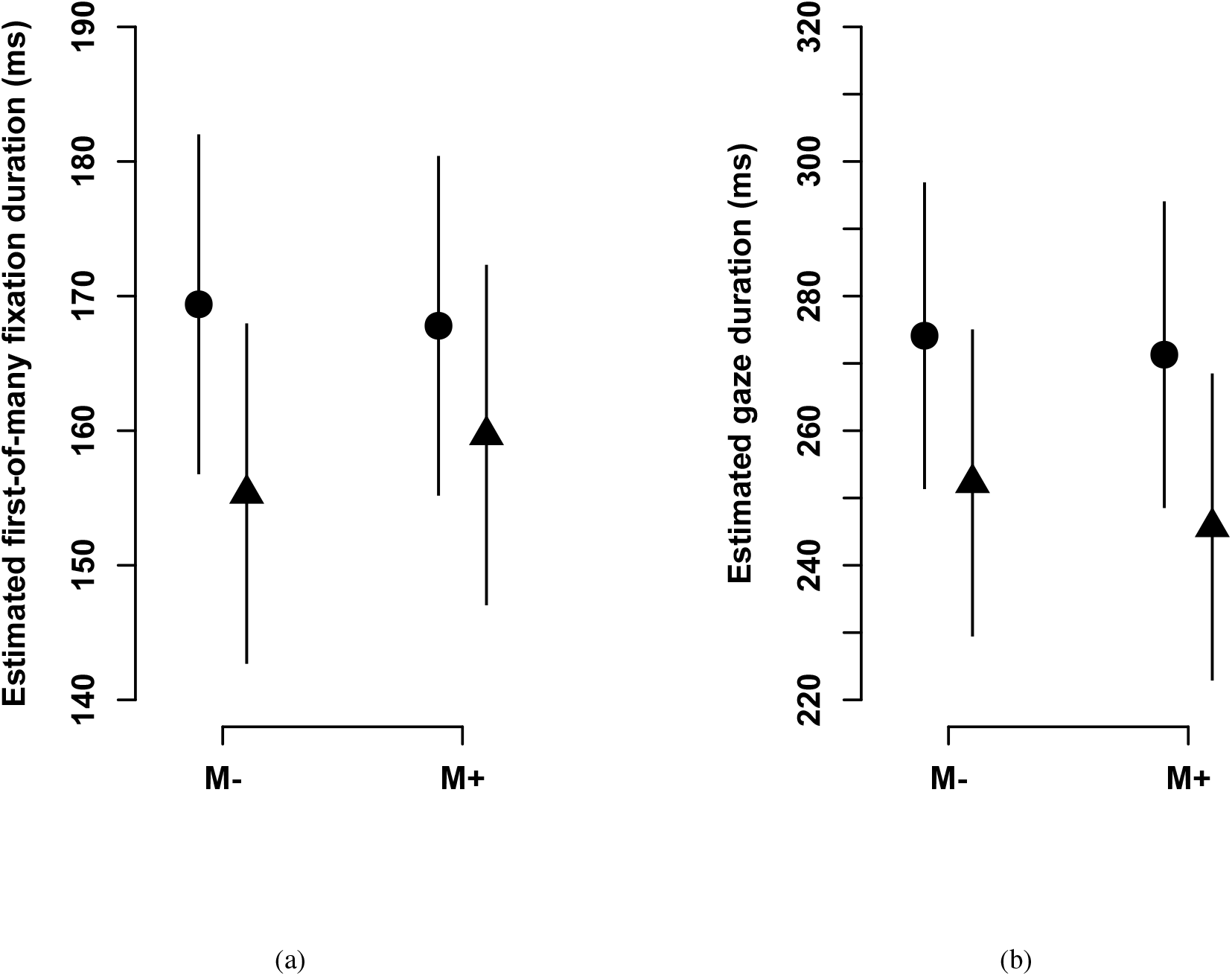
LMM estimates for first-of-many fixations duration (panel a) and gaze duration (panel b) on the target words. Error bars represent 95% CIs. Note: ●/▲, semantically incongruent/congruent; M-/+, morphologically incongruent/congruent.

Gaze duration (GD) revealed the same pattern. We observed solid semantic priming effect, *F*(1,3004) = 8.658, *p* = .004 (corresponding model parameter, *t*(3004) =2.943), with no morphological priming, *F*(1, 3004) = .337, *p* = .562, nor interaction between semantic and morphological congruity, *F*(1, 3004) = .053, *p* = .818. The estimated effect size was 23.742 ms for semantic priming and 4.670 ms for morphological priming. Figure 2b presents the model-based estimates for the four design cells. This pattern did not change when we considered only those trials in which the primes were fixated (semantic effect: *F*(1,2864) = 5.914, *p* = .015).

The same pattern emerged again in total looking time (TLT)—we observed solid semantic priming effect, *F*(1, 3004) =19.253, *p* < .001 (model parameter, *t*(3004) = 4.388), with no morphological priming *F*(1, 3004) = .709, *p* = .401, nor interaction between semantic and morphological congruity *F*(1,3004) = .239, *p* = .626. This pattern did not change when we considered only those trials in which the primes were fixated (semantic effect: *F*(1,2864), *p* =). The estimated effect size was 67.356 ms for semantic priming and 12.872 ms for morphological priming.

As anticipated in Section 2.2, we wanted to exclude the possibility that the small difference in predictability between S+ and S-conditions would be the main driver of the semantic effects. Therefore, we checked if semantic priming remains significant in models that excludes items with high CP values (>.10) and where, as a consequence, conditions are even more closely matched for predictability^6^. This was indeed the case for first-of-many fixations duration *F*(1, 2255) = 4.63, *p* = .033, gaze duration, *F*(1,2259) = 7.43, *p* = .007, and total looking time, *F*(1,2259) = 15.86, *p* < .001, while the effect could no longer be observed for target-on-prime, *F*(1, 2227) = .29, *p* = .591.

### 3.3 EEG results

The topography of the FRP grand averages for the semantic and morphological priming conditions are shown in Figure 3. Around 100ms after the onset of the target fixation, we can observe that all conditions (top and middle row of both panels in Figure 3) generated a positive-polarity response over the posterior areas of the scalp. This kind of response has previously been reported in other coregistration/natural reading studies, where it has been marked either as a P1 (Degno et al., 2018) or as a lambda response (Dimigen et al., 2012; Kazai and Yagi, 2003). Independently of the terminology, this response is equivalent to the P1 component in the classic ERP literature, and is thought to be visually-evoked. While this response was largest over the posterior areas of the scalp, it also surfaces in the frontal electrodes with a reversed polarity (see also Figure 4, where this response can be observed in the time series). Further, at around 400ms after fixation onset, the comparison between semantically related and unrelated primes (Figure 3a, bottom) revealed a dipolar response with positive polarity over the frontal electrodes and negative polarity over the posterior areas of the scalp. No such response is observed for the comparison of morphologically related and unrelated condition (Figure 3b, bottom).

**Figure 3:**
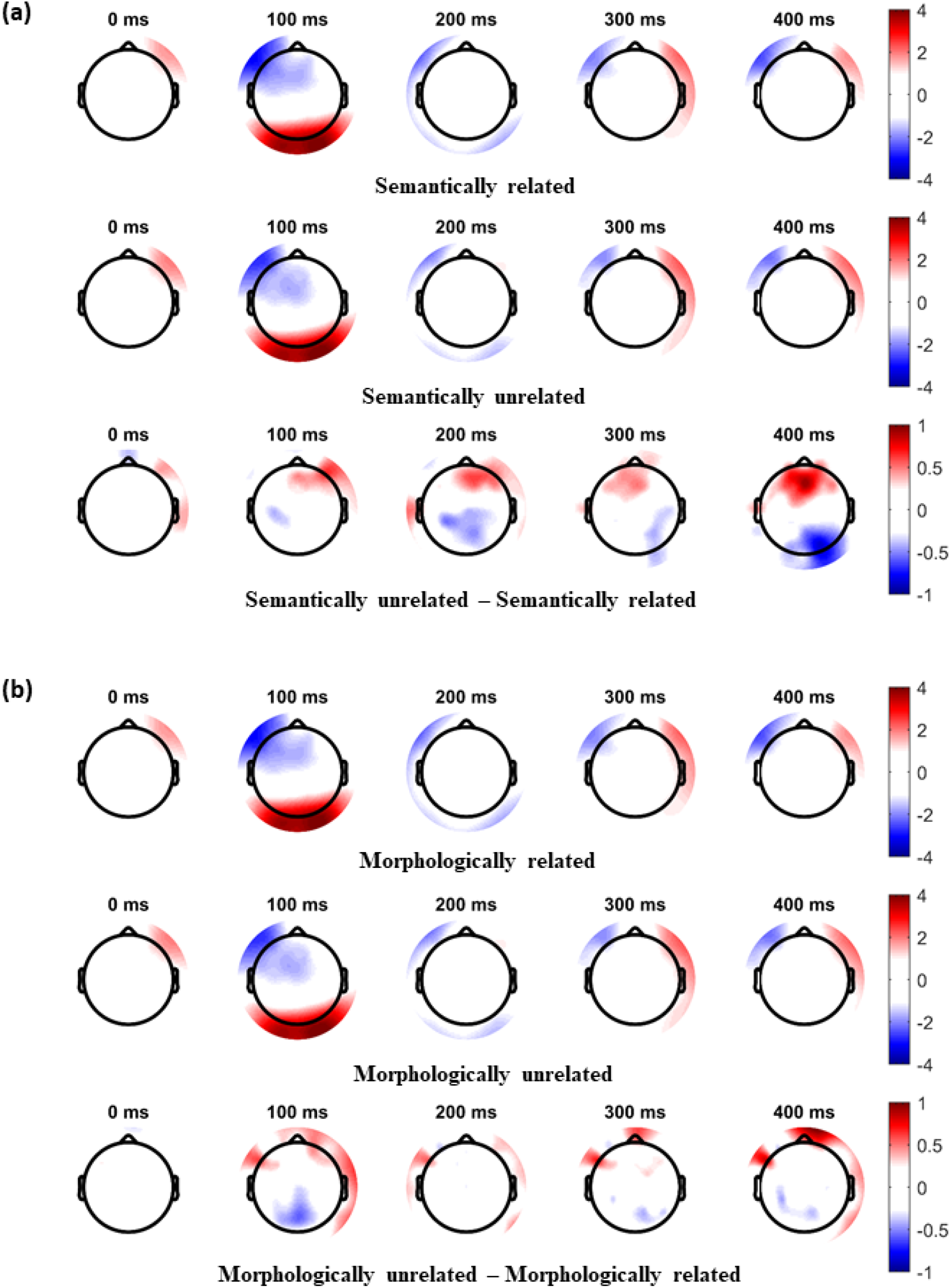
FRP results, time-locked to the fixation onset on the target word. Panel (a): Grand average FRPs in response to semantically related (top) and unrelated (middle) prime-target pairs, and the difference between the two (bottom). Panel (b): Grand average FRPs in response to morphologically related (top) and unrelated (middle) prime-target pairs, and the difference between the two (bottom).

**Figure 4:**
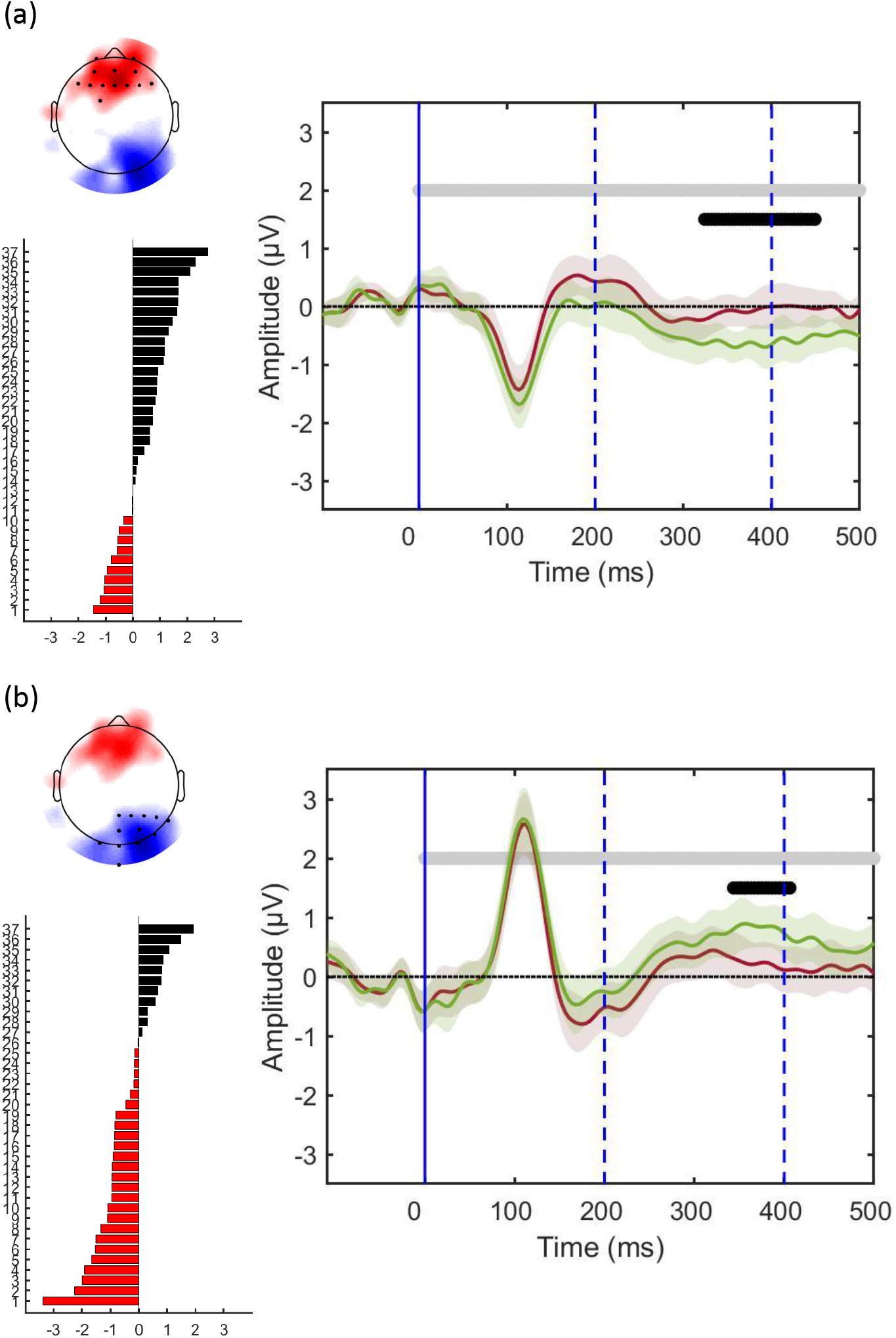
Grand average FRPs for the first (a) and the second cluster (b) emerging from the semantic comparison. FRPs are time-locked to the fixation onset on the target word (solid vertical line), in response to the semantically related (green) and unrelated (brown) prime word. Error bars denote 95% CI. The horizontal grey lines delimit the time window of interest, that is, the time points that were considered in the cluster analysis. The horizontal black lines denote the time windows of the observed cluster; the top left topoplot represents the cluster scalp distribution. The bottom left graph shows the strength of the statistical evidence for that cluster across participants.

Semantic priming was confirmed, and emerged as statistically solid, in the clustering analysis that compared the semantically related (S+) and semantically unrelated (S-) conditions, averaged across morphological congruency (M+M-). Unrelated primes elicited a frontal positive inflection (Figure 4a, top left) starting at around 190 ms, and reaching a peak between 300-450 ms after the fixation onset on the target word (Figure 4a, right). The observed cluster reached significance in the latter time window (*g* = −0.564, 95%*CI* = [.125, 1.004], *p* = .032). The single subject plot shows that the effect is quite consistent across participants (Figure 4a, bottom left).

A second cluster emerged in the semantic comparison over the right posterior electrodes, which likely reflects the negative part of the same dipole (Figure 4b, top left). The time pattern is very similar to the positive cluster: semantically related and unrelated primes start to diverge around 180 ms after the fixation onset, and become maximally different between 350ms and 400 ms (Figure 4b, right). This second cluster, however, does not reach statistical significance (*g* = −.539, 95%*CI* =[−.982, −.096], *p* = .125).

Similarly, we compared morphologically related (M+) to morphologically unrelated (M-) conditions, averaged across semantic congruency (S+S-). Unlike the semantic effects, morphological congruency does not show any statistically significant modulation of the signal: we only observed one candidate cluster which did not reach significance (*g* = −.591, 95%*CI* = [−1.02, −.166], *p* = .83).

Further, we assessed whether there was any modulation of the semantically related conditions by morphological congruency (i.e. S+M+ vs. S+M-) and, symmetrically, whether there was any modulation of the morphologically related conditions by semantic congruency (i.e., S+M+ vs. S-M+). No statistically significant effect was observed in this analysis (all p>.2), thus suggesting that semantic and morphological priming do not interact, similarly to what emerged in the eye tracking analysis.

## 4 Discussion

Building on similar eye tracking paradigms and taking advantage of some handy features of the Slovenian language, we analyzed eye movement behavior and Fixation-Related Potentials (FRP) time-locked on the fixation onset on the target word, in order to investigate semantic and morphological cross–word priming during natural sentence reading. Semantic facilitation emerged clearly, both in the oculomotor behavior and in FRP components. This effect was not modulated by morphological agreement, nor we observed any morphological priming in the first place. Importantly, all these results emerged in a very natural sentence reading paradigm, thus revealing lexical dynamics as they likely emerge in our everyday reading experience. Moreover, these data extends our theoretical knowledge by showing us what happens when the visual word identification system is engaged with several words in a short amount time.

Our eye movement results confirm previous observations of cross–word semantic priming in natural sentence reading (Carroll and Slowiaczek, 1986; Camblin et al., 2007). Importantly, the current experiment allows to rule out word predictability in the sentence context as a source for this priming. Despite target words being barely predictable, looking times were shorter on words when the preceding content word had a similar meaning. Also, semantic priming remained solid when we sub-set our data so as to provide even more stringent matching for predictability across conditions.

Existing models of isolated word identification and standard accounts of priming between individual words suggest a fairly straightforward interpretation, which relates to residual activation in the lexical-semantic system from word N-1 when a semantically related word N comes to the stage. This just seems the logical consequence of how the lexical-semantic system has been conceived for decades, as a network of interrelated nodes that interact with each other depending on the similarity of their representations (e.g., Hoffman et al., 2018; McRae and Boisvert; Meyer and Schvaneveldt, 1971). Also, semantic priming was shown to operate on a time scale that is largely compatible with the timing of fixation on two consecutive content words; in classic, single-word priming experiments, semantic effects hold firm when the Stimulus-Onset Asynchrony (SOA) between prime and target is in the range of the duration of a fixation (~200 ms; e.g., Nadalini et al., 2018; Rastle et al., 2000b).

However, if we consider individual word identification in the broader context of sentence processing, this interpretation becomes less incidental. In fact, the meaning of each individual word must integrate into the overall semantic message of the sentence, and this might well happen, at least in principle, by ‘merging’ the meaning of multiple words within the same lexical-semantic system that allows the identification and interpretation of each individual word. From this perspective, the brain would code for the meaning of, e.g., *red house* exactly by keeping the representations for *red* and *house* active within the lexical-semantic system at the same time^7^. So, it would not be by accident that activation holds in the system long enough for multiple words to be coded at the same time; this would be the core feature that allows us to understand phrases and sentences. It is interesting to note, then, that several recent models in the field of Computational Linguistics operate just on this assumption; the meaning of a sentence is the concatenation of the meaning of each individual word (e.g., Radford et al., 2019; Devlin et al., 2019; Vaswani et al., 2017). Of course, these models are not necessarily cognitive models (Guenther et al., 2019), but the fact that they are able to capture language production with unprecedented accuracy might mean that at least some of their computational assumptions are psychologically realistic.

Of course, this is not the only logical possibility. In fact, some models of sentence processing have indeed suggested an alternative approach, whereby lexical-semantic information on individual words is gathered in the lexicon, and then sent to a different cognitive structure for sentence-level integration. For example, in Snell et al. (2018)’s OB1, words that reach an identification threshold are selected to populate a ‘sentence-level representation in working memory’, where the meaning of the whole message is presumably computed. This alternative approach is not necessarily incompatible with cross-word semantic priming; in fact, activation coming from separate words is still likely to co-exist in the lexical-semantic system for long enough to generate priming. This would probably depend on the model specific parameters, which find in these novel data some further constraint that will help to fine-tune the theory. Also, it is not entirely clear what happens to a word’s activation in OB1, once that word reaches the identification threshold and gets selected for the sentence-level representation: does it get shut down? Or perhaps it decays progressively? This is also likely to impact the ability of OB1 to yield cross-word semantic priming. It is interesting to note, however, that the dynamics that would be responsible for cross-word priming do not carry any functional role here: it is essentially an ‘accident’ that the activation remains in the lexicon long enough to generate priming. In fact, one could in principle shut down this activation in a way that prevents cross-word priming, and the model would still be able to operate nicely, because the meaning of the sentence as a whole is computed outside of the lexicon. On the contrary, if the computation of the sentence meaning rests on the co-existence of activation from multiple words within the lexicon, cross-word priming would be intrinsic to a well-functioning system, and therefore it can never be killed without the system losing the ability to understand sentences. We leave this prediction as a nice challenge for future work, both computational and experimental (e.g., with neuropsychological patients who might, or might not show a dissociation between sentence comprehension and cross-word priming).

This eye tracking data are augmented by simultaneously recorded FRP data, time-locked to the fixation onset on the target word. The first notable component to mention here is a robust visually-evoked component, which emerged 100ms after fixation on the target word in all conditions as a positive-polarity response over the posterior region. Importantly, this result is in line with the previous natural reading coregistration studies (Dimigen et al., 2012; Degno et al., 2018), which have suggested that this component is an equivalent to the well-explored P1 in the ERP literature, where it is thought to reflect visual encoding (see, e.g., Dien (2008) for a review). With this, our data provides an important additional piece of evidence that this finding generalizes to natural reading.

Additionally, our data revealed an important novel finding—a modulation of a later FRP component by semantic relationship between the target word and its preceding prime. Compared to semantically related, semantically unrelated primes triggered a frontal positive-going component in the FRP, which started around 190ms and reached its maximum between 300ms and 400ms after fixation onset. In the same time window, a comparable effect, yet with reversed polarity, is observed over posterior brain areas, but doesn’t reach statistical significance. Since the two FRPs are acting in a very similar manner, we believe that the current data suggests them to be positive and negative part of a single dipole.

The timing of this effect makes it compatible with the classic N400 component for semantic priming, which is in line with the interpretation offered by previous FRP studies (Kretzschmar et al., 2009; Dimigen et al., 2012). However, while previous studies reported this modulation either in the reading of a list of words (i.e.,Dimigen et al., 2012) or in antonym sentence that introduced semantic violations (e.g., ‘ the opposite of black is white/yellow/nice’; Kretzschmar et al., 2009), here we observe it in the natural reading of perfectly well-formed sentences. This shows that the neural dynamics that tracks semantic priming between individual words, or in ill-formed sentence contexts, actually extend to a condition that is more akin to everyday-life reading.

Importantly, we observed N400/semantic priming in an environment that rules out word predictability as a possible source—our target words were not strongly constrained by the sentences, and semantic priming remained solid even when we excluded from the analyses the few target words with cloze probability higher than .10. This adds to a large literature investigating the nature of the N400 effect in individual word processing (e.g., Delaney-Busch et al., 2019; Bentin et al., 1985; Lau et al., 2013), or in sentence processing paradigms where words were delivered one by one (e.g. Wlotko and Federmeier, 2012; Kutas and Hillyard, 1984b). Without negating, of course, the sensitivity of N400 to word predicition, which is very well proven in these studies, here we show that this component can track semantic similarity as reflected in priming from semantic memory even when words are barely predictable at all within sentences. This sits well with the results of another FRP study, which showed independent N400 effects for predictability and spreading activation using non-sensical sentences (e.g., ‘ The opposite of black is *yellow’ Kretzschmar et al., 2009); here we extend these results to the natural reading of perfectly well-formed material.

In contrast, our data do not reveal any effect of morphological priming, either in the oculomotor behavior or in FRP components. These eye movement results nicely complement previous eye tracking experiments (e.g., Paterson et al.,2011)—when semantics and morphology are manipulated independently like in this experiment, the latter does not seem to give rise to cross-word processing savings during sentence reading. FRP components, instead, suggest that in natural reading, sharing an abstract morphological inflection, at least when denoted by distinct orthographic realizations, does not modulate any of the components traditionally linked with morphological processing, such as N250 and N400, as observed in ERP/RSVP paradigms (e.g., Lavric et al., 2010; Smolka et al., 2015; Rodriguez-Fornells et al., 2002; Rastle et al., 2015).

A note of caution is in order here, though. Especially in the individual word literature, morphological priming is typically addressed through shared stems or affixes that are fully realised orthographically (e.g., *dealer–deal, kindness–softness*; Crepaldi et al., 2016; Rastle et al., 2004; Marslen-Wilson, 2007). Here, instead, primes and targets shared an abstract morphological inflection, which was denoted by different affixal, orthographic realizations (e.g., ‘*avtomobil-a‘, (two) cars*, and *‘mačk-i‘, (two) cats*). This approach allows us to rule out any orthographic (or phonological) contribution to morphological effects, which is why we adopted it. However, it may also justify the discrepancy between what we find here and what is reported in the ERP/RSVP paradigms (e.g., Lavric et al., 2010; Smolka et al., 2015; Rodriguez-Fornells et al., 2002; Rastle et al., 2015), as well is in the individual word priming literature showing solid morphological facilitation (e.g., Marslen-Wilson et al., 1994; Feldman, 2000; Rastle et al.,2000a; Gonnerman et al., 2007).

Another important aspect of the results of the present study is the stark contrast between semantic and morphological processing during sentence reading. Not only semantic facilitation emerges while morphological priming does not, but also we were unable to see any interaction between the two players—the semantic effect was not affected by whether primes and targets were inflected alike, nor the morphological effect was modulated by semantic similarity. This may indicate that abstract morphological agreement is computed, at least in part, outside of the lexical-semantic system that gives rise to semantic priming; within this system, morphological facilitation would only emerge when the priming morpheme is fully expressed orthographically (e.g., Crepaldi et al., 2016; Duñabeitia et al., 2008). Also, this lack of interaction between semantic and morphological relatedness might support theories that suggest largely distinct lexical-semantic and morphological systems (e.g., Mcbride-Chang et al., 2008; Ramirez et al., 2014). Whatever theoretical interpretation one may want to adopt here, these data suggest that morphological processing is more locally encapsulated, in a way that prevent processing spillover between neighbouring words.

In addition to providing theoretical insight, the data described in the present paper open a few interesting questions, which the novel paradigm adopted here may help addressing.

First, an important next step would be to study cross-word priming with different types of similarities between primes and targets, such as case/gender agreement or orthographic similarity. This latter seems particularly interesting, as individual word priming suggests that orthographic overlap triggers lexical competition (e.g., Crepaldi et al., 2016; Davis and Lupker, 2006), which would raise a prediction for inhibitory cross-word priming between orthographically similar words. Such inhibitory priming effects in sentence reading have previously been reported in some studies (e.g., Paterson et al., 2009), while others seems to suggest that orthographic overlap between words embedded in sentences only triggers inhibitory priming when primes and targets are also phonologically related (e.g., *wings–kings*, vs. *bear–gear*; Frisson et al., 2014).

Another interesting issue is related to the distance between primes and targets. In our sentences, they were only separated by a short, high-frequency conjunction word; and always sat within the same coordinating phrase. How much lag is cross–word priming able to overcome? And how would syntax play out here? We have observed that morphological inflection does not seem to affect semantic facilitation during sentence reading; would it be the same for perhaps more prominent morpho–syntactic factors such as phrase boundaries, or word movement traces?

## Footnotes

1 Of course, different models will make different predictions as to which specific words will get activated, and under which specific dynamics. The point that we are trying to make here, however, generally applies to all models of visual word identification, so we ignore the differences between them and focus on what they have in common.

2 Of course, several other types of priming are reported in the literature. We won’t focus here on repetition priming (e.g., fence-FENCE) or nonword priming (e.g., bouse-HOUSE) for reasons that will become apparent when we get to our experimental design; we want to test priming within natural sentences, and word repetitions and nonwords very rarely/never appear within sentences. We might have considered lexical, orthographic priming (e.g., able-AXLE), but the nature of this type of priming is more controversial, both experimentally and theoretically (e.g., Davis, 2006). We will consider orthographic priming more in depth in the Discussion.

3 There were only 3 trials with a single fixation on the target word. Thus, the analysis of first-of-many fixation duration corresponds to an analysis of first fixation duration.

4 Number of trials (mean (SD), minimum number of trials) per condition for each participant: S+M+: 15.51 (2.52), 8; S+M-: 14.86 (2.47), 9; S-M+: 15.30 (2.86), 8; S-M-: 15.22 (2.29), 10

5 Please note that, because the experiment was not designed specifically for testing this effect, priming is assessed across different words here. Results should thus be taken with some caution.

6 With this procedure, we lost 746 items, which amounts to 25% of the dataset.

7 Of course, the meaning of word combinations is not always an entirely transparent juxtaposition of the meaning of the two words, for example in compounds and idiomatic expressions. There is a very rich literature on this issue (e.g., Flick et al., 2017; Marelli et al., 2017), which proposed several possible solutions for these cases, including independent representations in the lexicon for these complex items (e.g., Crepaldi et al.,2010; Grainger and Beyersmann, 2017; Taft, 2004)

## Notes

### Competing Interest Statement

The authors have declared no competing interest.

